# Obesity Potentiates T_H_2 Immunopathology via Dysregulation of PPAR*γ*

**DOI:** 10.1101/825836

**Authors:** Sagar P. Bapat, Yuqiong Liang, Sihao Liu, Ling-juan Zhang, Ian Vogel, Darryl J. Mar, Carmen Zhou, Eun Jung Choi, Christina Chang, Nanhai He, In-kyu Lee, Jae Myoung Suh, Laura E. Crotty Alexander, Christopher Liddle, Annette R. Atkins, Ruth T. Yu, Michael Downes, K. Mark Ansel, Alexander Marson, Richard L. Gallo, Ronald M. Evans, Ye Zheng

## Abstract

How obesity affects immune function is not well understood. Clinically, obesity is strongly associated with severe T_H_2 immunopathology^1-3^, though the physiological, cellular, and molecular underpinnings of this association remain obscure. Here, we demonstrate that obese mice are susceptible to severe atopic dermatitis (AD), a major manifestation of T_H_2 immunopathology and disease burden in humans^4,5^. Mechanistically, we show that dysregulation of the nuclear hormone receptor (NHR) PPAR*γ* (peroxisome proliferator-activated receptor gamma) in T cells is a causal link between obesity and the increased T_H_2 immunopathology. We find that PPAR*γ* oversees a cellular metabolic transcriptional program that restrains nuclear gene expression of the chief T_H_2 priming and effector cytokine interleukin-4 (IL-4). Accordingly, thiazolidinediones (TZDs), potent PPAR*γ* agonists, robustly protect obese mice from T_H_2 immunopathology. Collectively, these findings establish PPAR*γ* as a molecular link between obesity and T_H_2 immune homeostasis and identify TZDs as novel therapeutic candidates for T_H_2 immunopathology. Fundamentally, these findings demonstrate that shifting physiologic metabolic states can shape the tone of adaptive immune responses to modulate differential disease susceptibility.

---

Obesity is a metabolic state that is characterized by several pathophysiological processes, imposing severe disease burden in the world today^6-8^. The immune system plays a fundamental role in restraining and potentiating much of this disease burden; however, little is known about how obesity shapes the immune system to drive immunopathology. Recent epidemiological and clinical studies have uncovered associations between obesity and AD, a prominently T_H_2-driven immunopathology in humans^2,9^, and the striking co-increase in worldwide incidence of obesity and AD, suggest the two seemingly independent disease entities may be related^6,8,10^. To assess the relationship between obesity and AD, we challenged obese mice fed high-fat diet (HFD) with a well-characterized experimental model of MC903-induced AD on the ear^11,12^ (Figure 1a). Strikingly, the HFD mice, compared to control lean mice, displayed an overall increased inflammatory response, as measured grossly by a ∼3-fold increase in change in ear thickness (Figure 1b). Consistently, physical examination of the MC903-treated ears revealed prominent erythema and scale which was increased in the HFD mice, hallmarks of dermal inflammation (Figure 1c). Under histological evaluation, the obese mice exhibited increased epidermal acanthosis, hyperkeratosis, and spongiosis, hallmark characteristics of human AD. Impressively, the dermis was significantly expanded with leukocytic inflammation (dashed line), which was relatively diminished in the lean mice (Figure 1d).

**Figure 1.**
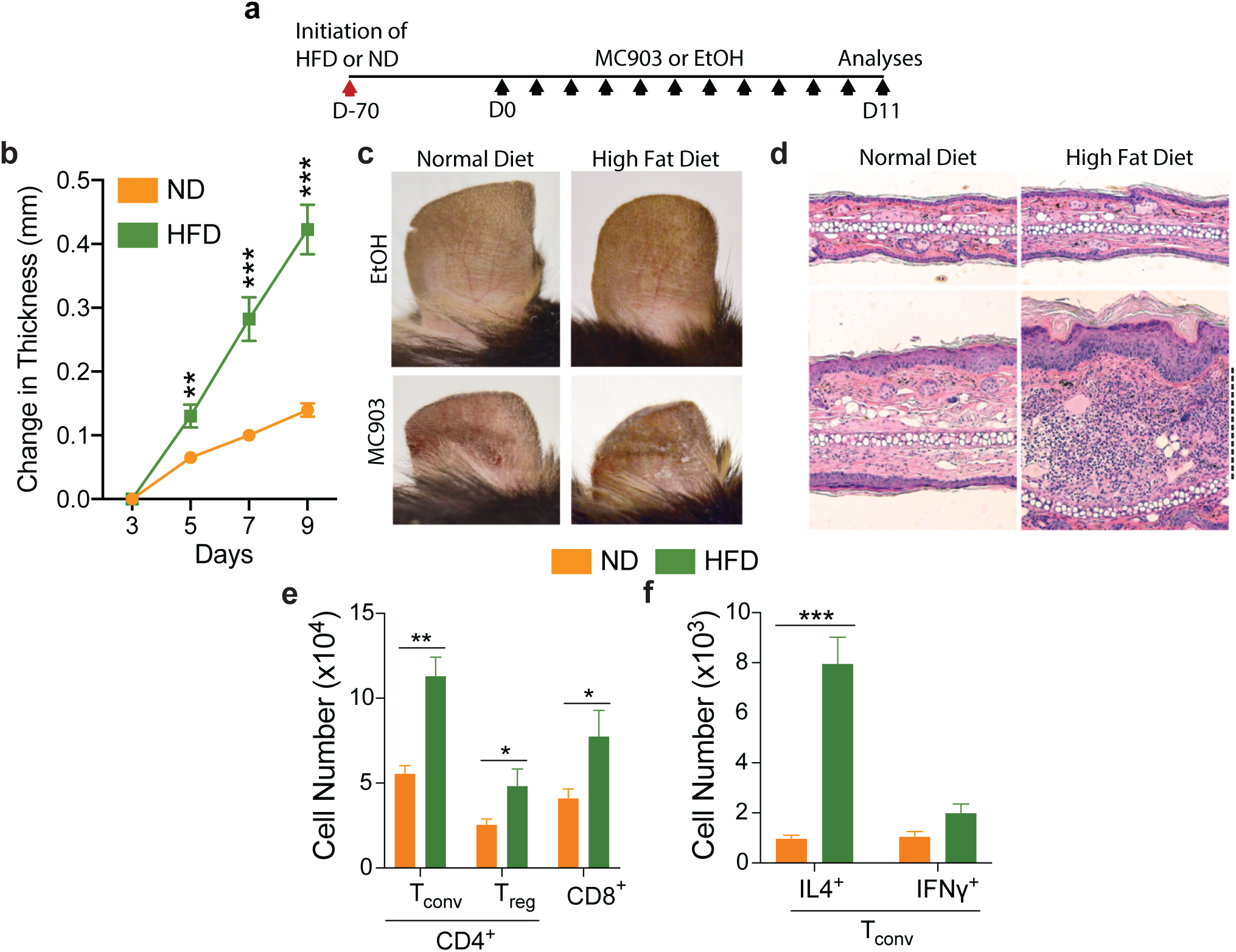
Obesity provokes increased T_H_2 immunopathology. **a,** Scheme of MC903-induced atopic dermatitis disease model used in this study. Black arrows indicate application of MC903 or EtOH administration to ear. **b,** Change in ear thickness during development of atopic dermatitis as indicated in **a. c,** Representative pictures of ears at Day 11. **d,** Representative images of hematoxylin and eosin stained histology of ears at Day 11. Scale bars, 100μm. **e,** Total T_conv_, T_reg_, and CD8^+^ T cell number from whole ear at Day 11. **f,** Total CD4^+^ IL-4^+^ and IFN ^+^ cell numbers from whole ear at Day 11. HFD, High fat diet; ND, Normal diet; D, Day. n=5 for ND and HFD for all comparisons. Data are mean ± s.e.m. \*\**P* < 0.01, \*\*\**P* < 0.001, Students *t-*test.

Since atopic disease is classically driven by T_H_2 immunity^13-15^, we sought to further characterize the infiltrating T cells induced by our experimental model of AD. Flow cytometric analyses revealed that the CD4^+^ conventional T cell (T_conv_), regulatory T cell (T_reg_), and CD8^+^ T cell populations were all increased in the obese mice (Figure 1e); however, the T_conv_ cells were the predominant T cell population and were significantly increased in their IL-4-competent (∼8.2 fold increase) but not IFN*γ*-competent populations compared to the lean mice (Figure 1f), an indicator of an increased T_H_2-polarized response. Of note, the link between obesity and T_H_2 immunopathology is dose-dependent as mice fed short-term HFD also display an increased inflammatory response upon induction of experimental AD, though to a lesser extent compared to the conventional HFD obese mice (Extended Data Figure 1a-f).

Nuclear hormone receptors (NHRs) are sensors of shifting physiological states^16,17^, and we wondered how NHRs were expressed in different types of T_H_ subsets and if any NHRs were specifically upregulated in T_H_2 cells. We surveyed the expression of NHRs in *in vitro* differentiated T_H_1, T_H_2, and T_H_17 cells and identified *Pparg* (and its heterodimeric partner *Rxra*) as highly and differentially expressed in T_H_2 cells (Extended Data Figure 2a). Further focused validation evaluating end-point and time-course gene expression as well as protein expression clearly identified PPAR*γ* as differentially and highly expressed in T_H_2 cells (Extended Data Figure 2b-d). Further, a differential analysis of genes involved in transcriptional regulation across *in vitro* differentiated T_H_1, T_H_2, and T_H_17 cells revealed *Pparg* expression to be highly specific to T_H_2 cells (Extended Data Figure 2e.) Consistent with our experiments, PPAR*γ* has recently been demonstrated as a transcription factor important for regulating transcriptional networks in T_H_2 cells, though its physiological relevance and functional importance is not well understood^18-20^.

PPAR*γ* and its ligands are master regulators of adipogenesis^21-24^. It is highly expressed in adipocytes and several other cell types (including immune cells such as fat-resident T_reg_ cells^25,26^) found within adipose tissue. Since obesity-associated dysregulation of PPAR*γ* activity in adipose tissue is an established cause of adipose tissue insulin resistance, we hypothesized that PPAR*γ* may be expressed *in vivo* in T_H_2 cells and similarly undergo obesity-associated dysregulation to promote a maladaptive T_H_2 immune responses in obese mice^22,24,27^. Indeed, by sorting on the infiltrating CD4^+^ T_conv_ and T_reg_ cells of lean mice, we were able to detect an appreciable 2-fold increase in *Pparg* in the T_conv_ but not T_reg_ cells compared to their respective splenic populations (Extended Data 2f).

These findings collectively motivated us to model T-cell specific PPAR*γ* dysregulation *in vivo* by creating T-cell specific PPAR*γ*-deficient mice (*Cd4*^*Cre*^ *x Pparg*^*fl/fl*^, PPAR*γ* TKO.) Importantly, T cell-specific loss of PPAR*γ* does not elicit any overt systemic inflammation associated with T cell dysregulation. PPAR*γ* TKO mice have normally sized spleens and lymph nodes (Extended Data Figure 3a,b) and equivalent splenic and lymph node T_reg_ (CD25^+^ Foxp3^+^) and activated (CD62L^lo^ CD44^hi^) CD4^+^ T cell populations compared to their control (*Cd4*^*Cre*^ *x Pparg*^*+/+*^) counterparts (Extended Data Figure 3c,d). Additionally, PPAR*γ* TKOs exhibit normal thymic T cell development as measured by CD4 and CD8 expression of thymic T cells (Extended Data Figure 3e). However, remarkably, when challenged with experimental AD under normal chow conditions, the PPAR*γ* TKO mice demonstrated a strikingly heightened T_H_2-driven immune response with a ∼2.7-fold increase in change in ear thickness and ∼60% increase in absolute ear thickness compared to control mice (Figure 2a,b). This heightened gross pathology was consistent with the histological examination which recapitulated many features of the wild-type obese mice including the most striking histologic feature – lymphocytic expansion of the dermis (Figure 2c, dashed line). Further flow cytometric analysis of the infiltrating immune cells demonstrated a ∼3-fold increase in IL-4-competent CD4^+^ T_conv_ cells (Figure 2d) with similar numbers of IFN*γ*-competent CD4^+^ T_conv_ cells (Figure 2e). By contrast, when obese PPAR*γ* TKO and controls were challenged with experimental AD, the differences in gross and histological pathology were largely diminished (Figure 2a-c). Additionally, flow cytometric analyses revealed no appreciable difference between obese PPAR*γ* TKO and control mice in IL-4-competent T_conv_ cells and IFN*γ*-competent T_conv_ cells (Figure 2d,e). These results suggest that HFD-induced obesity is largely epistatic to T cell-specific genetic deficiency of *Pparg* in controlling T_H_2 immune homeostasis, linking the obesity-associated susceptibility to T_H_2 immunopathology with dysregulation of PPAR*γ* in T cells.

**Figure 2.**
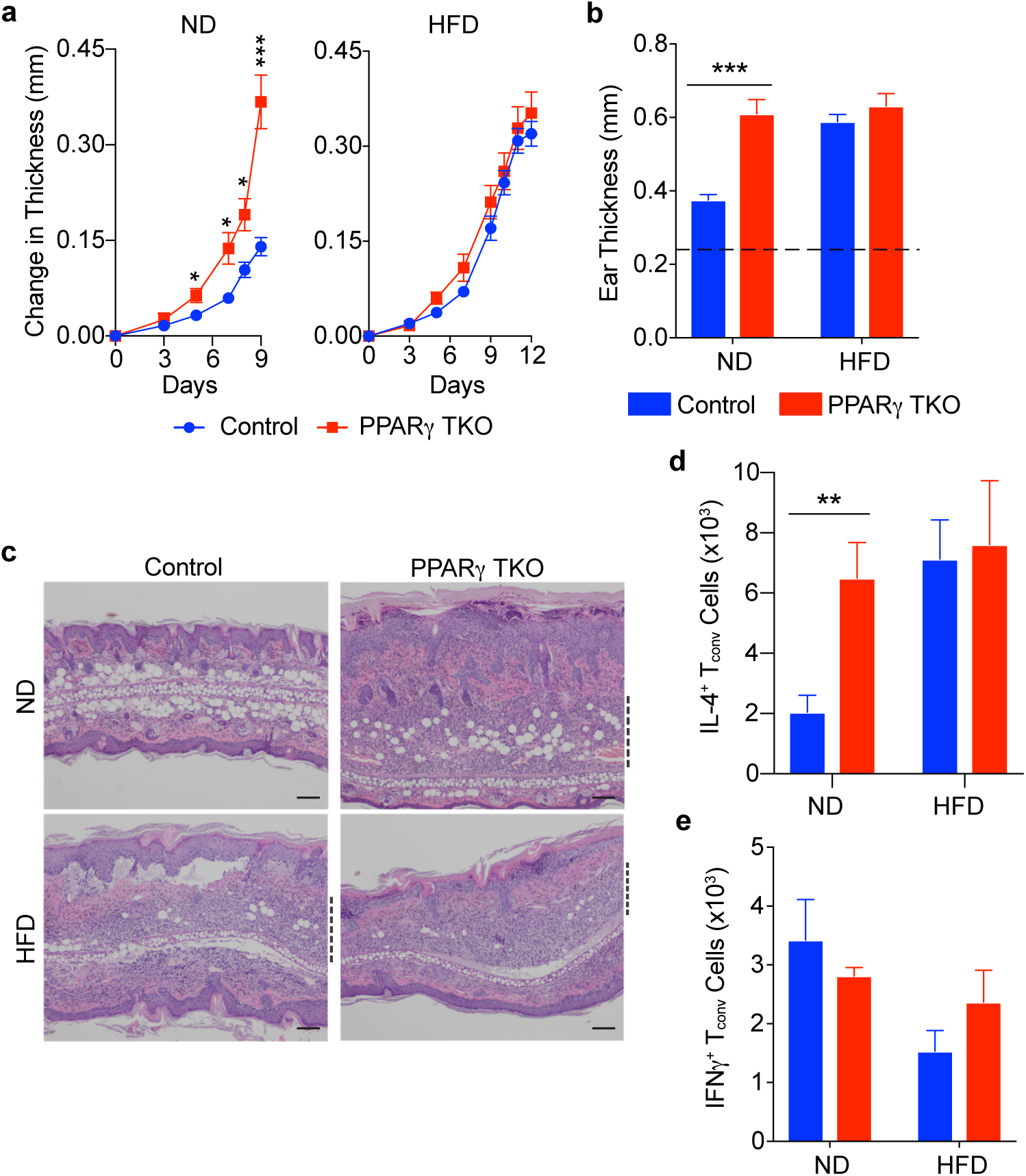
Obesity is epistatic to T cell-specific PPAR deficiency in potentiating atopic disease. **a,** Change in ear thickness during development of atopic dermatitis with mice fed ND or HFD. **b,** Absolute ear thickness at Day 11. Dotted line indicates average ear thickness at Day 0 of all mice in the study. **c,** Representative images of hematoxylin and eosin stained histology of ears at Day 11. Scale bars, 100μm. **d,e,** Total IL-4^+^ T_conv_ cells (**d**) and IFN ^+^ T_conv_ cells (**e**) from whole ear at Day 11. HFD, High fat diet; ND, Normal diet; Control, CD4^Cre^; PPAR TKO, CD4^Cre^ PPAR ^fl/fl^; n=5 for all groups in figure, except (**e**) where 3 PPAR TKO were on ND and 4 PPAR TKO were on HFD. Data are mean ± s.e.m. \**P* < 0.05, \*\**P* < 0.01, \*\*\**P* < 0.001, Student’s *t-*test.

Our *in vivo* studies revealed PPAR*γ* as a T cell-intrinsic restraint on T_H_2 immune responses. Although the transcriptomic and cistromic biology of PPAR*γ* has been studied extensively in the context of macrophages and dendritic cells^28-31^, the genomic regulation of PPAR*γ* in T cells is less studied. Intriguingly, a recent study utilizing genome-wide CRISPR knock-out screens included PPAR*γ* as part of the core transcriptional factors regulating T_H_2 activation and identity^20^; however, the functional role of PPAR*γ* activation or deficiency on T_H_2 cells has not been advanced. To gain mechanistic insights into how PPAR*γ* may be regulating T_H_2 immune responses and restraining T_H_2-driven immunopathology, we conducted transcriptomic analyses of *in vitro* differentiated T_H_2 cells from wild-type or PPAR*γ* TKO mice with or without treatment with the PPAR*γ*-agonist rosiglitazone (Rosi) to experimentally activate the PPAR*γ* transcriptional program. 1071 genes were differentially expressed, and hierarchical clustering revealed clear clusters of Rosi-responsive genes that were dependent on PPAR*γ* – the Rosi Up Cluster (455 genes) and Rosi Down Clusters I (171 genes) and II (361 genes) (Figure 3a). To complement transcriptomic analyses, we empirically identified the DNA binding sites for PPAR*γ* in T_H_2 cells via ChIP-Seq. The top DNA-binding motif discovered was the canonical PPAR-response element (PPRE, Figure 3b.) 2534 binding sites mapping to 1563 genes were identified, of which 215 genes were members of the Rosi Up or Down I and II clusters (Figure 3c). Unexpectedly, ontological analyses of these 215 genes identified only one consensus GO term: response to organic cyclic compounds – a parent term for a large group of metabolism-related ontologies (Figure 3d). In contrast, when similar analyses were applied to the entirety of the genes found in the Rosi Up and Rosi Down clusters, the ontologies were exclusively immunological for the Rosi Down Clusters and a mix of metabolic and immunological for the Rosi Up Cluster (Figure 3d). Taken together, this suggests that PPAR*γ* activation directly activates genes involved in cellular metabolism whose collective expression and function then alters the expression of the genes associated with immunological processes. Congruent with this model, several metabolism genes markedly induced by Rosi treatment exhibited high PPAR*γ* ChIP peak scores, whereas several key immunological genes induced by Rosi treatment had largely diminished or absent PPAR*γ* ChIP peak scores (Figure 3e). Importantly, *Il4,* the chief T_H_2 priming and effector cytokine, was found in the Rosi Down II Cluster and had an absent PPAR*γ* ChIP peak score (Figure 3f), as was the case with all cytokines found in the Rosi Down Clusters. This Rosi-induced, PPAR*γ*-dependent decrease in *Il4* was further confirmed by analysis of targeted gene expression, intracellular staining, and secretion (Extended Data Figure 4a-c).

**Figure 3.**
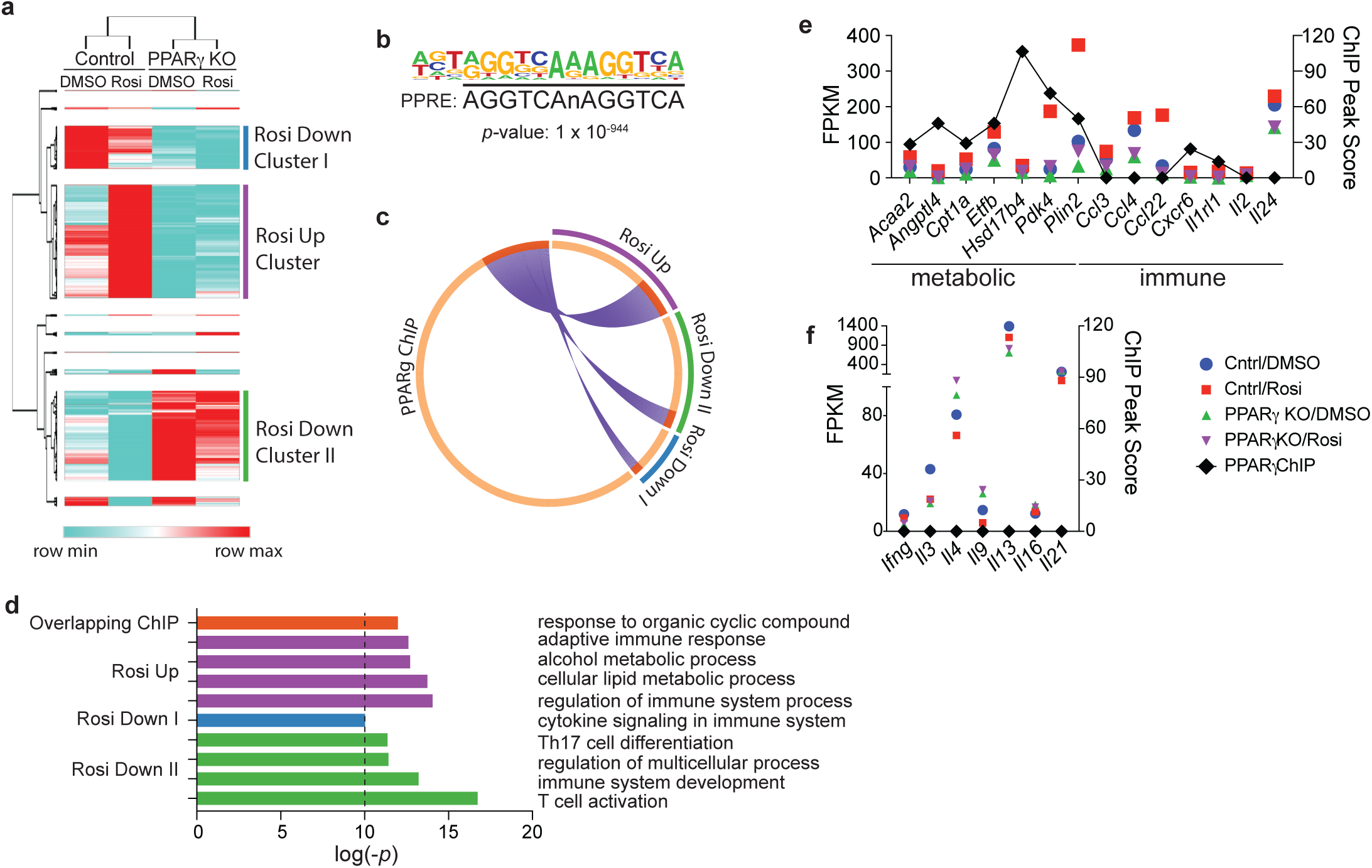
PPAR directly controls T_H_2 cellular metabolism. **a,** Hierarchical clustering of differentially expressed genes between T_H_2 cells sufficient or deficient in *Pparg* treated with rosiglitazone (Rosi) or dimethyl sulfoxide (DMSO) (cells pooled from 4 mice, same data set used in **c-f**). **b,** Top scoring DNA motif of PPAR ChIP-Seq peaks in T_H_2 cells via de novo analysis (cells pooled from 4 mice, same data set used in **c-f**). **c,** Circos plot representing gene lists from the Rosi Up, Rosi Down I, and Rosi Down II Clusters along with the peak lists from the PPAR ChIP-Seq of T_H_2 cells. Purple arcs link identical gene names from two different lists. **d,** Bar graph representing P values of gene ontology clusters with P<10^−10^ of indicated gene lists. Each cluster is labeled with the most representative gene ontology term for that cluster. **e,f,** Selected FPKM and corresponding PPAR ChIP peak values of genes from the Rosi Up cluster (**e**) and Rosi Down clusters (**f**) that are metabolic or immunological in function.

Metabolic regulation of immune function occurs in a highly context and cell-type specific manner^32-34^. To further understand the mechanism by which PPAR*γ*-activated metabolic reprogramming could control immunological function in T_H_2 cells, we examined genes in the Rosi Up Cluster that were also direct targets of PPAR*γ*. We found genes whose consensus activities would promote *de novo* lipogenesis (*Plin2, Gpam, Dgat1*), fatty acid oxidation (*Pdk4, Cpt1a, Acaa2, Eci2, Slc25a20, Acadl, Acsl5*), and mitochondrial mass (*Etfb, Mrpl45, Mrpl1*), which would likely manifest in mitochondrial substrate usage of fatty acids (Extended Data Figure 4d,e). Indeed, follow-up studies showed a clear Rosi-induced, PPAR*γ*-dependent increase in mitochondrial mass (Extended Data Figure 4f-h) as well as mitochondrial oxidation of fatty acids (Extended Data Figure 4i,j) with no increase in cellular glucose uptake (Extended Data Figure 4k) or in mitochondrial oxidation of glucose-derived carbons (Extended Data Figure 4l). Collectively, these studies argue for PPAR*γ* as a core T_H_2 transcription factor enforcing mitochondrial oxidative capacity required to restrain over-exuberant T_H_2 effector function.

Due to the robust ability of Rosi to decrease T_H_2 effector function and enforce lipid catabolic pathways in a PPAR*γ*-dependent fashion *in vitro*, we hypothesized that perhaps Rosi treatment could be used to protect obese mice from exacerbated AD. Remarkably, mice treated with Rosi had substantially reduced disease, as measured by a ∼40% reduction in increase in ear thickness (Figure 4a,b), marked reduction in lymphocytic infiltration (Figure 4c), and a significant decrease in IL-4 competent T_conv_ cells (Figure 4d), effects that were largely dependent on T cell-specific PPAR*γ*. Interestingly, we did observe effects of Rosi that were not dependent on T cell-specific PPAR*γ*, including a modest increase in disease severity (Figure 4a-c) and a reduction in IFN*γ*-competent T_conv_ cells (Figure 4e), possibly related to T cell-extrinsic mechanisms of PPAR*γ* action, which has been shown to be expressed in several skin cell types like adipocytes, sebocytes, hair follicles, and keratinocytes^35^.

**Figure 4.**
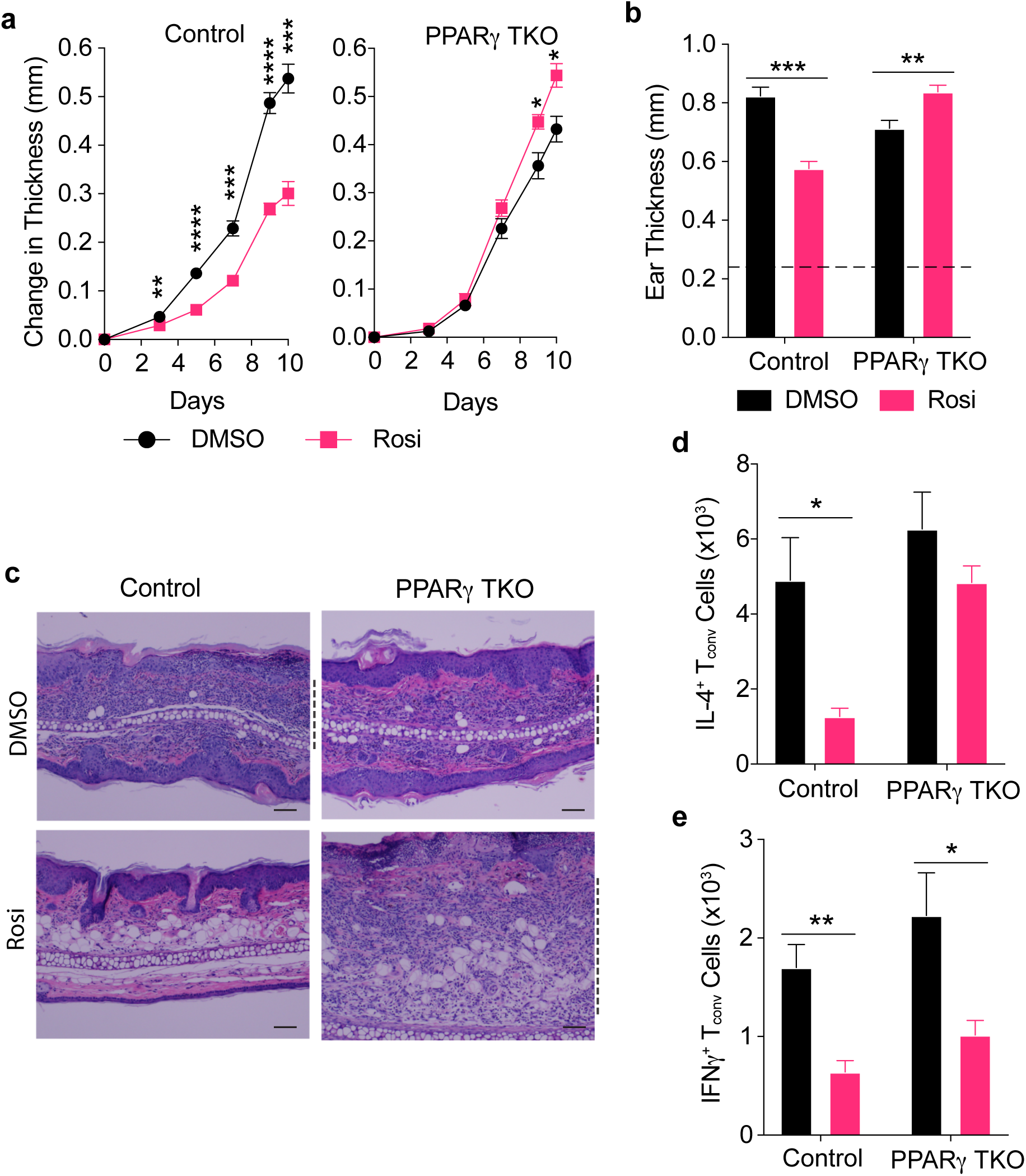
Pharmacological activation of PPAR protects against obesity-associated T_H_2 immunopathology in a T cell PPARγ-dependent manner. **a,** Change in ear thickness during development of atopic dermatitis with mice fed HFD diet or HFD with Rosi. **b,** Absolute ear thickness at Day 11. Dotted line indicates average ear thickness at Day 0 of all mice in the study. **c,** Representative images of hematoxylin and eosin stained histology of ears at Day 11. Scale bars, 100μm. **d,e,** Total IL-4^+^ T_conv_ cells (**d**) and IFN ^+^ T_conv_ cells (**e**) from whole ear at Day 11. PPAR TKO, CD4^Cre^ PPAR ^fl/fl^; n=5 for all groups, except (**e**) where 4 control mice were treated with HFD diet or HFD with Rosi. Data are mean ± s.e.m. \**P* < 0.05, \*\**P* < 0.01, \*\*\**P* < 0.001, \*\*\*\**P* < 0.0001, Student’s *t-*test.

Taken together, our studies provide a physiological, cellular, and molecular basis for the clinically observed association between obesity and T_H_2 immunopathology, establishing obesity-associated dysregulation of PPARγ in T cells as an important driver of obesity-associated T_H_2 immunopathology. Leveraging mouse genetics, we identify loss-of-function of PPARγ specifically in T cells as a key mechanistic step in the potentiation of T_H_2 immunopathology in the obese state. Unexpectedly, we find that PPARγ serves as a brake on effector function by controlling mitochondrial metabolism. Further, we demonstrate that the PPARγ agonists, TZDs, commonly prescribed as insulin-sensitizers for treatment of Type-2 diabetes, could be used to treat T_H_2 immunopathology, a novel indication. Fundamentally, we demonstrate that distinct physiological metabolic states can alter cellular metabolic states to shape immune responses in specific ways to modulate disease susceptibility.

## Acknowledgements

We would like to thank Y. Dai, J. Alvarez, A. Cheng, Y. Zhang for technical assistance, L. Ong and C. Brondos for administrative assistance, and A. Levine and S. Kajimura for critical review of the manuscript. S.P.B. was supported by U.S. National Institutes of Health (NIH) grants F30 DK096828 and T32 GM007198. L.Z. and R.L.G. are supported by NIH grants R01AR069653, U19 AI117673, R37 AI052453, and R01 AR076082. I.-K.L. is supported by grants of the Korea Health Technology R&D Project through the Korea Health Industry Development Institute (KHIDI), funded by the Ministry of Health & Welfare, Republic of Korea (HI16C1501) and Basic Science Research Program through the National Research Foundation (NRF) of Korea (NRF-2017R1A2B3006406). J.M.S. is supported by the National Research Foundation of Korea grants NRF-2018R1A2A3075389, NRF-2016M3A9B6902871. L.E.C.A. is supported by an American Heart Association grant 16BGIA27790079 and a VA BLR&D Career Development Award 1IK2BX001313. C.L. and M.D. are funded by grants from the National Health and Medical Research Council of Australia Project grants 512354, 632886 and 1043199. K.M.A. is supported by U.S. National Institutes of Health (HL107202, HL109102) and the Sandler Asthma Basic Research Center. A.M. holds a Career Award for Medical Scientists from the Burroughs Wellcome Fund, is an investigator at the Chan Zuckerberg Biohub, has received funding from the Innovative Genomics Institute (IGI), and is a member of the Parker Institute for Cancer Immunotherapy (PICI). R.M.E. is an Investigator of the Howard Hughes Medical Institute (HHMI) at the Salk Institute and March of Dimes Chair in Molecular and Developmental Biology, and is supported by U.S. NIH grants DK057978, HL088093, HL105278 and ES010337, the Leona M. and Harry B. Helmsley Charitable Trust 2017PG-MED001, Ipsen/Biomeasure, California Institute for Regenerative Medicine CIRM DISC2-11175, and The Ellison Medical Foundation. Y. Z. is supported by the NOMIS Foundation, the Rita Allen Foundation, and U.S. National Institutes of Health grants AI107027, CA014195, OD023689. This work was also supported by National Cancer Institute funded Salk Institute Cancer Center core facilities (CA014195) and the James B. Pendleton Charitable Trust. Research reported in this publication was supported by the National Institute of Environmental Health Sciences of the National Institutes of Health under Award Number P42ES010337. The content is solely the responsibility of the authors and does not necessarily represent the official views of the National Institutes of Health.

## Author Contributions

S.P.B. conceived of the research. S.P.B., R.M.E., and Y. Z. designed and supervised the research. S.P.B., Y.L., S.L., L.-J.Z., I.V., D.M., C.Z., E. J. C., C.C., N.H., L.E.C.A., and Y.Z. performed experiments. S.P.B., Y.L., S.L., J.M.S., I.-K.L., L.E.C.A., C.L., A.R.A., R.T.Y., M.D., R.M.E. and Y.Z. analyzed data. S.P.B. wrote the manuscript. S.P.B., A.R.A., R.T.Y., M.D., R.M.E. and Y.Z. reviewed and edited the manuscript.

## Author Information

RNA-Seq and ChIP-Seq data can be accessed in the NCBI Sequence Read Archive under the accession PRJNA553761. Reprints and permissions information is available at www.nature.com/reprints. Correspondence and requests for materials should be addressed to R.M.E. or Y.Z.

## Materials and Methods

### Mice

All mice were housed in the specific pathogen-free facilities at The Salk Institute for Biological Studies or purchased from The Jackson Laboratory. PPARγ TKO mice were generated by crossing CD4Cre^1^ transgenic mice and *Pparg*^fl/fl^ (ref. 2) mice. We used the *Foxp3*^Thy1.1^ (ref. 3) reporter mice when isolating T_reg_ and T_conv_ CD4^+^ cells from spleen and ear for subsequent RNA analysis. Mice within The Salk Institute for Biological Studies received autoclaved normal chow (MI laboratory rodent diet 5001, Harlan Teklad), irradiated HFD (60 kcal% fat, Research Diets), irradiated HFD mixed with rosiglitazone (15 mg kg^-1^ of food, Research Diets) or DMSO (Research Diets), or ND (10 kcal% fat, Research Diets). All mice used for studies were male. All procedures involving animals were performed in accordance with protocols approved by the Institutional Animal Care and Use Committee (IACUC) and Animal Resources Department (ARD) of the Salk Institute for Biological Studies.

### MC903-induced experimental atopic dermatitis

Mice were anesthetized via isoflurane and MC903 solution (0.1 mM in ethanol, 10 μL/ear, R&D Systems) was applied to mouse ears daily for 9-12 days. Ear thickness was assessed utilizing a micrometer (Mitutoyo, #227-211). Upon harvest, ears were dissected from the mice and prepared for histological analyses or flow cytometry.

### Single-cell suspension of ear-infiltrating immune cells

Dissected ears were minced into fine pieces (1-2 mm^3^), and digested in a stromal vascular isolation buffer (HBSS with Calcium and Magnesium, 20mg/mL BSA, 20mg/mL penicillin, 20mg/mL streptomycin) containing 2 mg ml^-1^ Collagenase D (Roche) at 37°C with intermittent shaking for 2 hrs. The suspension was then passed through a 100-μm mesh to remove undigested clumps and debris. The flow-through was centrifuged at 400 RCF for 10 min. The pellet containing the stromal vascular fraction was washed once in 10 mL RPMI, and the resultant isolated cells were subjected to FACS analysis directly or first stimulated with PMA and ionomycin in the presence of brefeldin A (GolgiPlug; BD) for 5 hrs at 37 °C for subsequent intracellular cytokine staining and FACS analysis.

### Histological analyses

Sections (4 μm) of fixed tissues were stained with haematoxylin and eosin according to standard procedures. Histopathological analyses were conducted on blinded samples for severity and extent of inflammation and morphological changes by a pathologist.

### Flow cytometry

The following antibodies were used. Biolegend: CD4 (RM4-5), CD8 (53-6.7), CD25 (7D4), CD44 (IM7), CD45.2 (104), CD62L (MEL-14), and IFN-γ (XMG1.2); eBioscience: Foxp3 (FJK-16s), IL-4 (11B11), IL-13 (eBio13A), TCRβ (H57–597). For intracellular staining, cells were treated with fixation and permeabilization reagents from BD or eBioscience (for Foxp3 staining) and labeled with appropriate antibodies before being analyzed on a BD FACSAria Cell Sorter. Data were analyzed using the BD FACSAria instrument (Becton Dickinson) and FlowJo software (FlowJo LLC).

### *In vitro* CD4^+^ T cell differentiation

CD4^+^ T cells were isolated from the spleen and lymph nodes using the EasySep Mouse CD4 Positive Selection Kit II (Stemcell Technologies). Naïve (CD25^-^CD44^hi^ CD62L^lo^) CD4^+^ T cells were sorted by flow cytometry from the bead purified CD4^+^ T cells. The naïve CD4^+^ T cells were resuspended in Click’s medium (Irvine Scientific) at 1 million cells per mL, and then plated on Day 0 in 24 well plates coated with goat-hamster IgG antibody (200ng/ml; MP Biomedicals) with the addition of soluble anti-CD3 (1μg/ml; 145-2C11) and anti-CD28 (1μg/ml; 37.51) from Bio X Cell. Polarizing conditions for different T helper subsets are as following: T_H_1: hIL2 (100U/ml; PeproTech), mIL-12 (20ng/ml; PeproTech) and anti-IL-4 (5μg/ml; Bio X Cell); T_H_2: hIL2 (100U/ml; PeproTech), mIL-4 (20ng/ml; Biolegend), anti-IFN-γ and anti-IL-12 (5μg/ml; Bio X Cell); T_H_17: mIL-6 (20ng/ml; Biolegend), hTGF-β (2ng/ml; PeproTech), anti-IFN-γ and anti-IL-12 (5μg/ml; Bio X Cell). When indicated, Rosi was dissolved in DMSO and added to a final concentration of 10μM on Day 0 and Day 3 of 4-5 day cultures.

### Primers for qPCR

*Il4* – For: AGGAGCCATATCCACGGATGCGA, Rev: TGTTCTTCGTTGCTGTGAGGACGT;

*Il13* – For: CACAGAAGACCAGACTCCCC, Rev: GTTGGTCAGGGAATCCAGGG;

*Pparg* – For: CACAATGCCATCAGGTTTGGG, Rev: GAAATGCTTTGCCAGGGCTC;

*Gata3* – For: CTTCCCACCCAGCAGCCTGC, Rev: CGGTACCATCTCGCCGCCAC;

*Tbx21* – For: GTCGCGCTCAGCAACCACCT, Rev: CGGCCACGGTGAAGGACAGG;

*Rorc* – For: CCGGACATCTCGGGAGCTGC, Rev: CGGCGGAAGAAGCCCTTGCA;

*Gapdh* – For: CAAGGTCATCCATGACAACTT, Rev: GGCCATCCACAGTCTTCTGG;

*Hprt* – For: GTCATGCCGACCCGCAGTCC, Rev: GGCCACAATGTGATGGCCTCCC;

*CoI* – For: TGCTAGCCGCAGGCATTAC, Rev: GGGTGCCCAAAGAATCAGAAC;

*Ndufv1* – For: CTTCCCCACTGGCCTCAAG, Rev: CCAAAACCCAGTGATCCAGC.

### Antibodies for western blot

Rabbit anti-PPARγ mAb (81B8, Cell Signaling), mouse anti-NDUFB8 mAb (20E9DH10C12, Invitrogen), mouse anti-Tubulin (DM1A, Sigma).

### ChIP-Seq library generation

Naïve CD4^+^ T cells were activated and polarized in T_H_2 conditions. On day 1 and 2, T cells were transduced with a retroviral vector expressing TY1-tagged *Pparg*. On day 4, transduced T cells were harvested for ChIP as described previously^4^ utilizing a TY1 antibody (Sigma, SAB4800032). ChIP-sequencing libraries were constructed and sequenced (50 bp single end reads) as described previously^4^. Short DNA reads were demultiplexed using Illumina CASAVA v1.8.2. Reads were aligned against the mouse mm10 reference genome using the Bowtie2 aligner with standard parameters that allow up to 2 mismatches per read. Peak calling, motif analyses, and other data analysis were performed using HOMER, a software suite for ChIP-seq analysis as described previously^4^. Visualization of ChIP-Seq results was achieved by uploading custom tracks onto the UCSC genome browser.

### RNA-Seq library generation and sequencing analysis

Total RNA was extracted from T_H_2 cells ∼96 hours after initiation of *in vitro* differentiation. RNA-sequencing libraries were prepared from 100 ng total RNA (TrueSeq v2, Illumina) and single-end sequencing was performed on the Illumina HiSeq 2500, using bar-coded multiplexing and a 100 bp read length, yielding a median of 34.1M reads per sample. Read alignment and junction finding was accomplished using STAR^5^ and differential gene expression with Cuffdiff 2^6^, utilizing UCSC mm10 as the reference sequence. Transcript expression was calculated as gene-level relative abundance in fragments per kilobase of transcript per million mapped fragments and employed correction for transcript abundance bias^7^. Metascape^8^ was utilized for functional annotation of genes. Morpheus (https://software.broadinstitute.org/morpheus) was employed for hierarchical clustering. Circos (https://circos.ca) was employed to make Circos plots^9^.

### IL-4 ELISA

Naïve CD4^+^ T cells were differentiated *in vitro* in T_H_2 polarizing conditions as described above in 24 well plates. 96 hours after initial activation, cells were collected, washed twice with PBS, and resuspended in T_H_2 polarizing medium without IL-4 in 24 well plates. The cells were cultured for an additional 48 hours and then the conditioned media was collected to quantitate IL-4 content by ELISA (BioLegend, Mouse IL-4 ELISA MAX Standard Sets) using absorbance at 490nm as a readout.

### Metabolic phenotyping via extracellular flux analysis

Mitochondrial substrate dependency and maximal respiration levels were determined by assessing oxygen consumption rate (OCR). OCR was measured using a 96 well extracellular flux analyser (Seahorse Bioscience). In brief, dead cells were removed from T cell cultures using Ficoll-Paque Premium 1.084 (17-5446-02; GE). 4 × 10^5^ live cells per well (generally 5 wells per sample) were spun onto a Cell Tak (Corning)-coated Seahorse plate and rested for 1 hour in a CO_2_ free incubator at 37 °C before commencing measurements using the Seahorse instrument. As indicated in figures, the following drugs were added into the wells (concentration is final concentration in wells): Oligomycin A (10 μM, Sigma), FCCP (5 μM, Sigma), Rotenone (7.5 μM, Sigma), Antimycin A (7.5 μM, Sigma), Etomoxir (100 μM, Sigma), UK5099 (2 μM, Sigma).

### Fatty acid oxidation assay

PPARγ TKO and control naïve CD4^+^ T cells were activated and polarized in T_H_2 conditions for 5 days. Rosi or DMSO was added to a final concentration of 10 μM on Day 0 and Day 3. On day 5, 5 million live cells were resuspended directly in 1 mL of loading media (DMEM, low glucose containing 2% fatty-acid free BSA (Sigma A6003), 0,25 mM carnitine (Sigma C-0158), and 2 μCi/reaction ^3^H-palmitic acid (American Radiolabeled Chemicals, Inc, ART 129)). Reaction mix was loaded onto a 24 well plate (200 μL per well). The cells were incubated for 2 hours and supernatant was collected from each well. The cells were harvested and washed with PBS three times and lysed in 0.1N NaOH to determine protein concentration for normalization. 100 μL of supernatants were treated with 100 μL of 10% trichloracetic acid (TCA). After mixing and 15 min incubation at room temperature, samples were centrifuged at maximum speed for 10 min. The resultant supernatant was transferred to a new microtube and treated with 100 μL of 5% TCA and 40 μL of BSA (10% BSA in TE buffer). After mixing and 15 min incubation at room temperature, samples were centrifuged at maximum speed for 10 min. ∼300 μL of resulting supernatant was transferred to a new microtube and 750 μL chloroform:methanol (2:1 mix) and 300 μL KCl:HCl (2M each, 3.6g KCl in 20 mL water, 4 mL conc. HCl) were added to the reaction. After mixing and 15 min incubation at room temperature, samples were centrifuged at maximum speed for 10 min. ∼600 μL of the resulting supernatant was collected and added to scintillation counting tubes containing 5 mL Ecolume (MP Biomedicals). Samples were mixed and subjected to measurement of radiation via scintillation counter (^3^H program).

### Glucose uptake assay

PPARγ TKO and control naïve CD4^+^ T cells were activated and polarized in T_H_2 conditions for 5 days. Rosi or DMSO was added to a final concentration of 10 μM on Day 0 and Day 3. 5 million cells per sample were resuspended in 1 ml of KRBH (Krebs-Ringer bicarbonate HEPES) buffer at pH of 7.4 (120 mM NaCl; 4 mM KH_2_PO_4_; 1 mM MgSO_4_; 0.75 mM CaCl_2_; 30 mM HEPES; 10 mM NaHCO_3_) and equally distributed into 5 wells of a 24 well plate. Cells were incubated for 30 min in 37 °C incubator, and then treated with 50 μL/well of loading buffer (0.5mM 2-DG (2-Deoxy-D-glucose) and 0.5Ci (3H-2DG)). Negative control wells were additionally treated with 10 μL/well of 1.5 mM cytochalasin B in DMSO to suppress active glucose uptake and control for non-specific uptake and binding of ^3^H-2-DG. Cells were incubated for 1 hour at 37 °C and then washed three times with PBS. Cells were lysed in 300 μL of 0.1 N NaOH. 200 μL of cell lysis/sample were collected for radioactivity measurements via scintillation counter. The remaining amount was utilized to conduct the Bradford assay to quantify protein concentration. All radioactivity counts were normalized to sample protein concentration, and the negative control counts were further subtracted from experimental sample measurements.

### Statistical analyses

Statistical analyses were performed with Prism 8 (GraphPad). *P* values were calculated using two-tailed unpaired or paired Student’s *t*-test. Mice cohort size was designed to be sufficient to enable statistical significance to be accurately determined. When applicable, mice were randomly assigned to treatment or control groups. No animals were excluded from the statistical analysis, with the exception of exclusions due to technical errors, and the investigators were not blinded in the studies. Appropriate statistical analyses were applied, assuming a normal sample distribution, as specified in the figure legends. No estimate of variance was made between each group. All *in vivo* experiments were conducted with at least two independent cohorts. All *in vitro* T_H_ differentiation experiments were conducted at least three times. ChIP-Seq and RNA-Seq experiments were conducted using multiple biological samples (as indicated in figure legends).

---

**Extended Data Figure 1.**
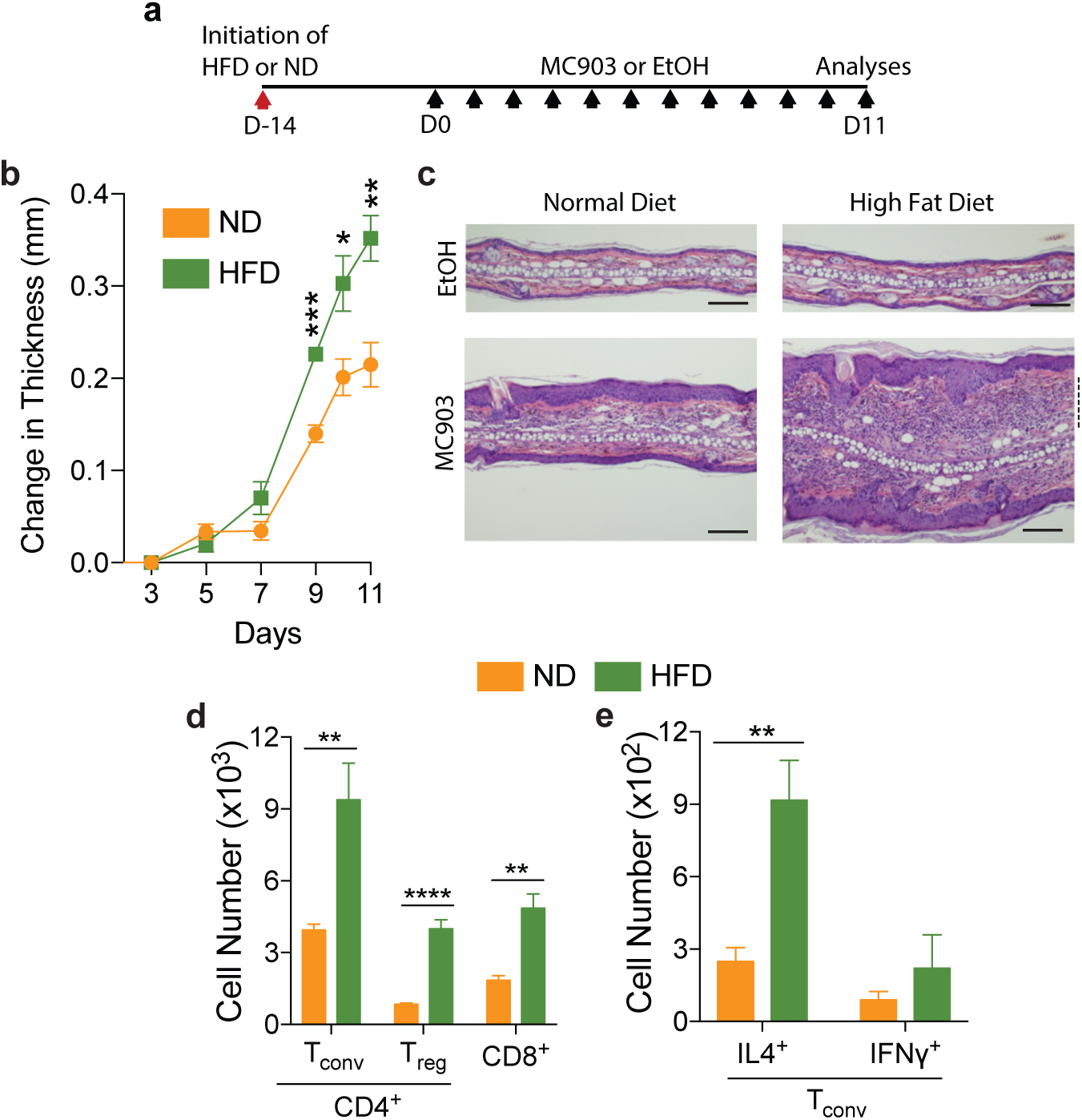
Short-term HFD provokes modestly increased T_H_2 immunopathology, though muted compared to long-term HFD. **a,** Scheme of MC903-induced atopic dermatitis disease model used in this study. Black arrows indicate application of MC903 or EtOH administration to ear. **b,** Change in ear thickness during development of atopic dermatitis as indicated in **a. c,** Representative images of hematoxylin and eosin stained histology of ears at Day 11. Scale bars, 200μm. **d,** Total T_conv_, T_reg_, and CD8^+^ T cell number from whole ear at Day 11. **e,** Total CD4^+^ IL-4^+^ and IFN ^+^ cell numbers from whole ear at Day 11. HFD, High fat diet; ND, Normal diet; D, Day. n=5 for ND and HFD for all comparisons. Data are mean ± s.e.m. \**P* < 0.05, \*\**P* < 0.01, \*\*\**P* < 0.001, Student’s *t-*test.

**Extended Data Figure 2.**
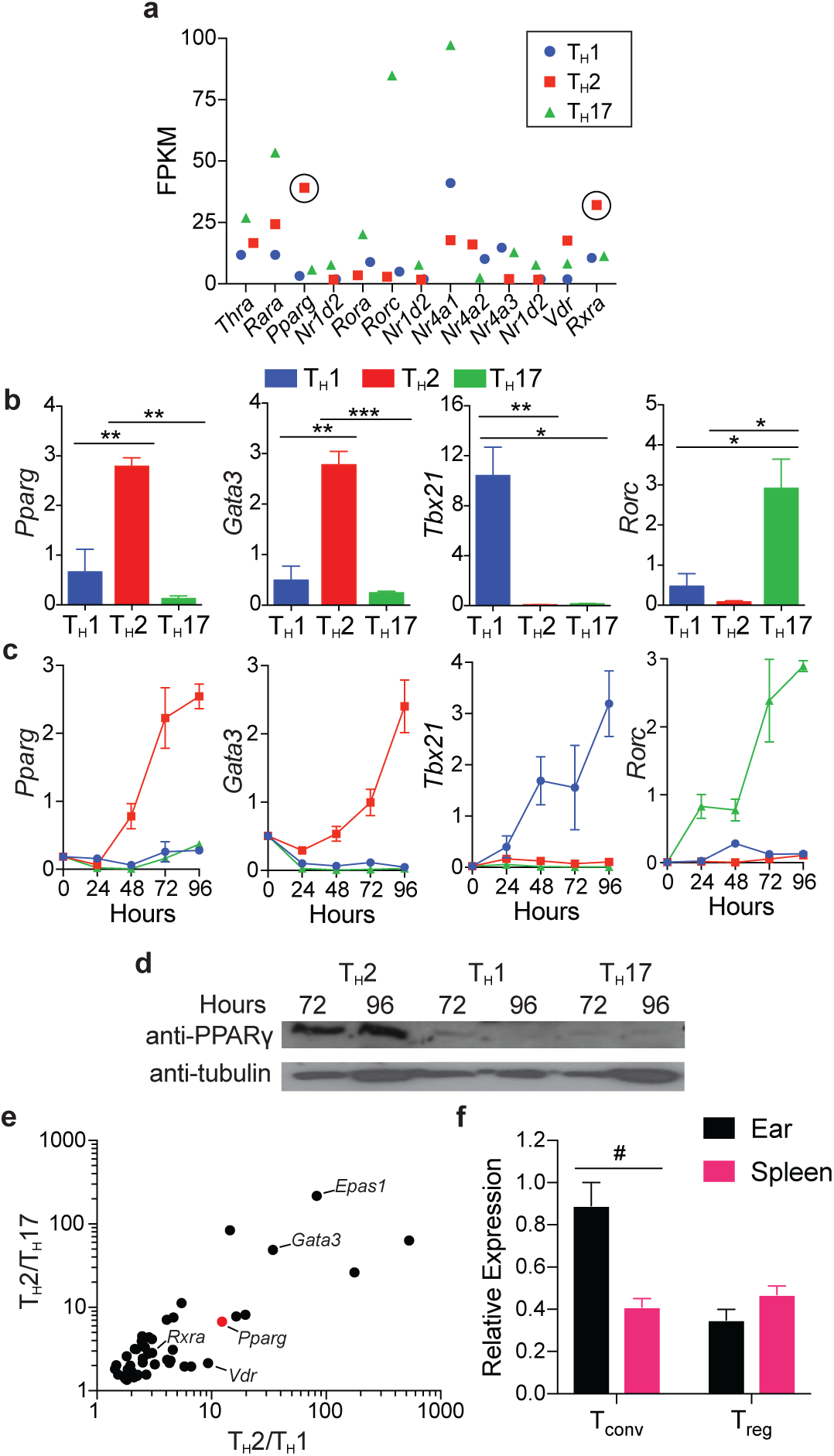
PPAR is differentially expressed in *in vitro* differentiated T_H_2 cells and in MC903-challenged infiltrating T_conv_ cells *in vivo*. **a,** Fragments per kilobase of transcripts per million mapped reads (FPKM) values of nuclear hormone receptor (NHR) superfamily genes differentially expressed in *in vitro* differentiated T_H_1, T_H_2, and T_H_17 cells. NHR genes that are differentially expressed in T_H_2 cells are encircled (cells pooled from 4 mice before inducing differentiation in triplicate, same data set used in **e**). **b,c** Relative expression (using *Hprt* expression as housekeeping gene) of indicated genes in *in vitro* differentiated T_H_1, T_H_2, and T_H_17 cells (cells pooled from 4 mice before inducing differentiation in triplicate) **b,** Gene expression determined at Hour 120 post induction of differentiation. **c,** Time course of gene expression from Hours 0-96 post induction of differentiation. **d,** Western blot of PPAR and tubulin at Hours 72 and 96 in *in vitro* differentiated T_H_1, T_H_2, and T_H_17 cells (cells pooled from 3 mice before inducing differentiation) **e,** FPKM values of genes that are differentially expressed in T_H_2 cells and involved in transcriptional regulation. Position of *Pparg* is marked with a red dot. **f,** *Foxp3*^Thy1.1^ mice were challenged with MC903-induced atopic dermatitis and T_reg_ and T_conv_ cells were sorted from spleen and ear and assayed for *Pparg* expression, relative to *Gapdh* (cells pooled from 4 mice in biological duplicates). Data are mean ± s.e.m. \**P* < 0.05, \*\**P* < 0.01, \*\*\**P* < 0.001, #*P*=.055, Student’s *t*-test.

**Extended Data Figure 3.**
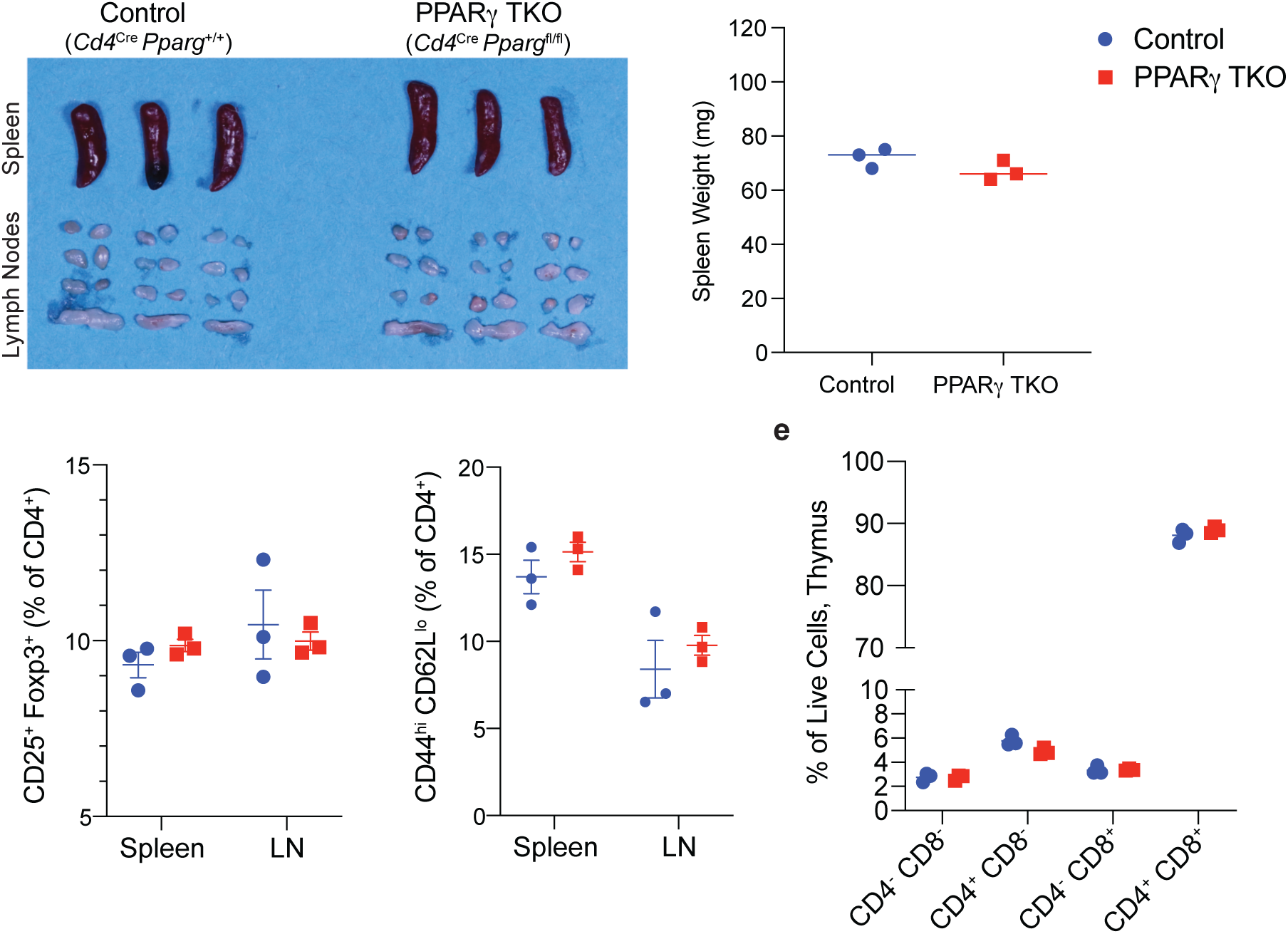
PPAR TKO mice display neither overt systemic inflammation nor altered T cell development. **a,** Picture of control and PPAR TKO spleens and lymph nodes. **b,** Spleen weights. **c,d,** T_reg_ (**c**) and activated T_conv_ (**d**) cell frequency in spleen and LN. **e,** Developing T cell subsets in thymus. n=3 mice per group. LN, lymph node. Student’s *t*-test.

**Extended Data Figure 4.**
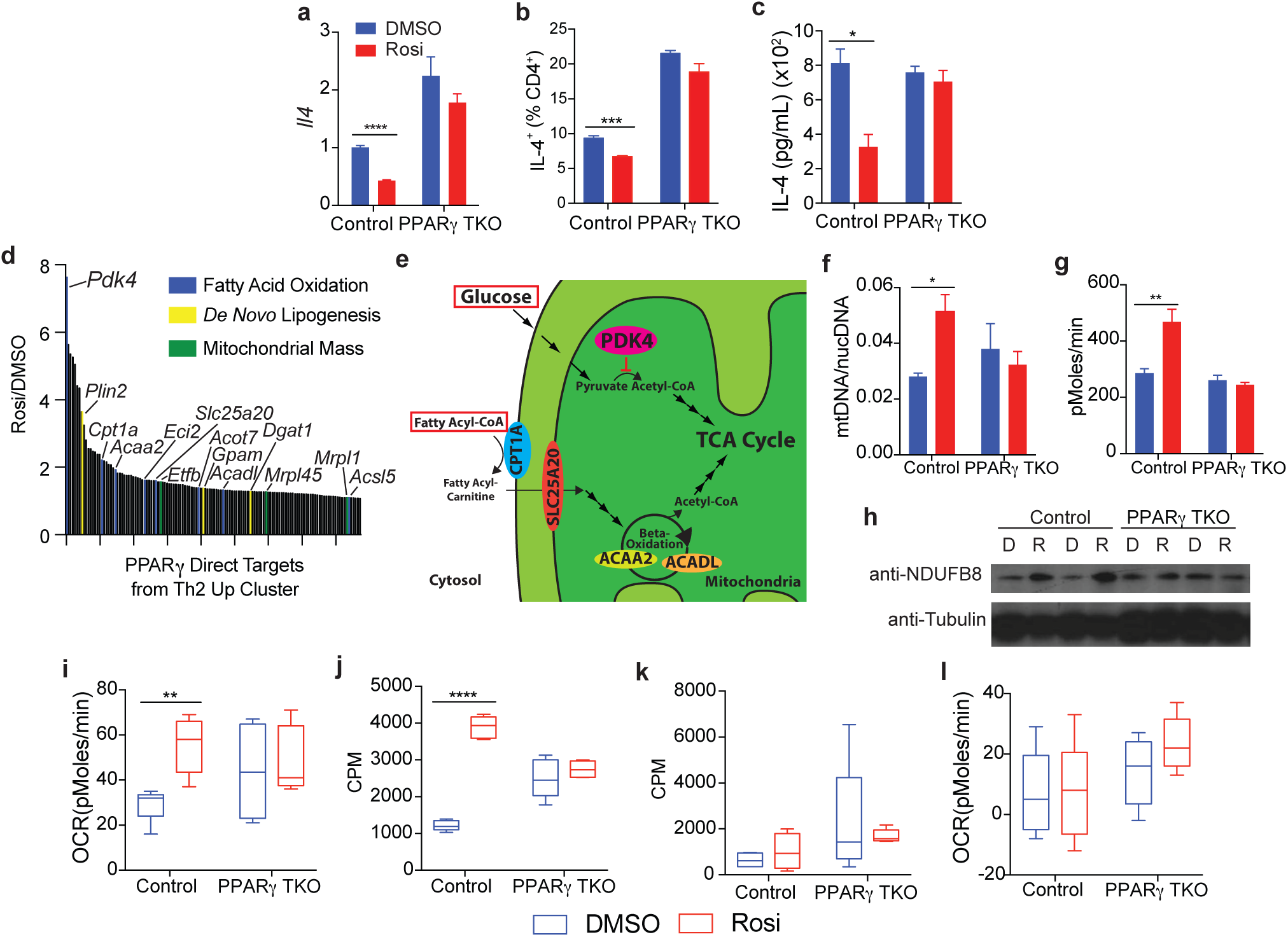
PPAR restrains IL-4 expression and drives a fatty acid catabolic program in T_H_2 cells. **a,** Relative expression (using *Hprt* expression as housekeeping gene) of *Il4* (n=3 per group). **b,c**, Intracellular protein expression via flow cytometry (**b**) and secretion via ELISA from conditioned media (**c**) of IL-4 (n=3 per group). **d**, PPAR direct targets as identified by ChIP that are upregulated upon PPAR activation (found in the Rosi Up clusters.) **e,** Graphical representation of the phenotypic change in the mitochondrial metabolism of T_H_2 cells upon PPAR activation. **f**, Mitochondrial DNA (*CoI*) to nuclear DNA (*Ndufv1*) ratio (n=3 per group). **g,** Maximal respiration after treatment with carbonyl cyanide-4 (trifluoromethoxy) phenylhydrazone (FCCP) (n=3 per group). **h,** Western blot of NDUFB8 (protein located in inner mitochondrial membrane) and tubulin at 96 hours in *in vitro* differentiated T_H_2 cells (cells pooled from 3 mice before inducing differentiation; D, DMSO; R, Rosi, 10μM). **i,l**, Mitochondrial substrate dependency of fatty acids (**i**) and glucose (**l**) measured by OCR after treatment with etomoxir and UK5099, respectively (OCR, oxygen consumption rate; n=5 per group). (**j,l**) Rates of beta-oxidation of exogenous fatty acids (**j**) and glucose uptake (**l**) (CPM, counts per minute, normalized to sample protein concentration; n=5 per group). All experiments performed with *in vitro* differentiated T_H_2 cells with cells pooled from 4 mice before inducing differentiation. Data are mean ± s.e.m. For box and whisker plots, whiskers plot min to max. \**P* < 0.05, \*\**P* < 0.01, \*\*\*\**P* < 0.0001, Student’s *t-*test.

## References

1. Hersoug, L. G. & Linneberg, A. The link between the epidemics of obesity and allergic diseases: does obesity induce decreased immune tolerance? Allergy 62, 1205–1213 (2007).

2. Zhang, A. & Silverberg, J. I. Association of atopic dermatitis with being overweight and obese: A systematic review and meta-analysis. J Am Acad Dermatol 72, 606–616.e4 (2015).

3. Fahy, J. V. Type 2 inflammation in asthma — present in most, absent in many. Nat Rev Immunol 15, 57–65 (2015).

4. Bieber, T. Atopic Dermatitis. N. Engl. J. Med. 358, 1483–1494 (2008).

5. Eckert, L. et al. The burden of atopic dermatitis in US adults: Health care resource utilization data from the 2013 National Health and Wellness Survey. 1–9 (2017). doi:10.1016/j.jaad.2017.08.002

6. The GBD 2015 Obesity Collaborators. Health Effects of Overweight and Obesity in 195 Countries over 25 Years. N. Engl. J. Med. 377, 13–27 (2017).

7. NCD-RisC, N. R. F. C. Articles Trends in adult body-mass index in 200 countries from 1975 to 2014: a pooled analysis of 1698 population-based measurement studies with 19·2 million participants. The Lancet 387, 1377–1396 (2016).

8. NCD-RisC. Worldwide trends in body-mass index, underweight, overweight, and obesity from 1975 to 2016: a pooled analysis of 2416 population-based measurement studies in 128·9 million children, adolescents, and adults. The Lancet 390, 2627–2642 (2017).

9. Beuther, D. A. & Sutherland, E. R. Overweight, Obesity, and Incident Asthma. Am J Respir Crit Care Med 175, 661–666 (2007).

10. Nutten, S. Atopic dermatitis: global epidemiology and risk factors. Ann Nutr Metab 66 Suppl 1, 8–16 (2015).

11. Li, M. et al. Topical vitamin D3 and low-calcemic analogs induce thymic stromal lymphopoietin in mouse keratinocytes and trigger an atopic dermatitis. PNAS 103, 11736–11741 (2006).

12. Leyva-Castillo, J. M. et al. Skin thymic stromal lymphopoietin initiates Th2 responses through an orchestrated immune cascade. Nature Communications 4, 1–12 (2015).

13. Mowen, K. A. & Glimcher, L. Signaling pathways in Th2 development. Immunol. Rev. 202, 203–222 (2004).

14. Locksley, R. M. Asthma and Allergic Inflammation. Cell 140, 777–783 (2010).

15. Bacher, P. et al. Regulatory T Cell Specificity Directs Tolerance versus Allergy against Aeroantigens in Humans. Cell 167, 1067–1069.e16 (2016).

16. Evans, R. M. & Mangelsdorf, D. J. Nuclear Receptors, RXR, and the Big Bang. Cell 157, 255–266 (2014). doi:10.1016/j.cell.2014.03.012

17. Lazar, M. A. Maturing of the nuclear receptor family. J Clin Invest 127, 1123–1125 (2017).

18. Chen, T. et al. PPAR-γ promotes type 2 immune responses in allergy and nematode infection. Sci Immunol 2, 1–11 (2017).

19. Nobs, S. P. et al. PPARγ in dendritic cells and T cells drives pathogenic type-2 effector responses in lung inflammation. J Exp Med 214, 3015–3035 (2017).

20. Henriksson, J. et al. Genome-wide CRISPR Screens in T Helper Cells Reveal Pervasive Crosstalk between Activation and Differentiation. Cell 176, 882–896.e18 (2019).

21. Forman, B. M. et al. 15-Deoxy-Δ12,14-Prostaglandin J_2_ Is a Ligand for the Adipocyte Determination Factor PPARy. Cell 83, 803–812 (1995).

22. Spiegelman, B. PPARγ: a nuclear regulator of metabolism, differentiation, and cell growth. J Biol Chem 276, 37731–37734 (2001).

23. Tontonoz, P. & Spiegelman, B. M. Fat and Beyond: The Diverse Biology of PPARγ. Annu Rev Biochem 77, 289–312 (2008).

24. Ahmadian, M. et al. PPARγ signaling and metabolism: the good, the bad and the future. Nat Med 19, 557–566 (2013). doi:10.1038/nm.3159

25. Bapat, S. P. et al. Depletion of fat-resident Treg cells prevents age-associated insulin resistance. Nature 528, 137–141 (2015). doi:10.1038/nature16151

26. Li, C. et al. TCR Transgenic Mice Reveal Stepwise, Multi-site Acquisition of the Distinctive Fat-Treg Phenotype. Cell 174, 285–299 (2018). doi:10.1016/j.cell.2018.05.004

27. Dubois, V. et al. Distinct but complementary contributions of PPAR isotypes to energy homeostasis. J Clin Invest 127, 1202–1214 (2017).

28. Chawla, A. et al. PPAR-gamma dependent and independent effects on macrophage-gene expression in lipid metabolism and inflammation. Nat Med 7, 48–52 (2017).

29. Odegaard, J. I. et al. Macrophage-specific PPARgamma controls alternative activation and improves insulin resistance. Nature 447, 1116–1120 (2007).

30. Szanto, A. et al. STAT6 transcription factor is a facilitator of the nuclear receptor PPARγ-regulated gene expression in macrophages and dendritic cells. Immunity 33, 699–712 (2010).

31. Daniel, B. et al. The Nuclear Receptor PPARγ Controls Progressive Macrophage Polarization as a Ligand-Insensitive Epigenomic Ratchet of Transcriptional Memory. Immunity 49, 615–626.e6 (2018).

32. Pollizzi, K. N. & Powell, J. D. Integrating canonical and metabolic signalling programmes in the regulation of T cell responses. Nat Rev Immunol 14, 435–446 (2014). doi:10.1038/nri3701

33. O’Neill, L. A. J., Kishton, R. J. & Rathmell, J. A guide to immunometabolism for immunologists. Nat Rev Immunol 16, 553–565 (2016).

34. Klein Geltink, R. I., Kyle, R. L. & Pearce, E. L. Unraveling the Complex Interplay Between T Cell Metabolism and Function. Annu Rev Immunol 36, 461–488 (2018).

35. Ramot, Y. et al. The role of PPAR γ-mediated signalling in skin biology and pathology: new targets and opportunities for clinical dermatology. Exp Dermatol 24, 245–251 (2015).

## Methods References

1. Lee, P. et al. A Critical Role for Dnmt1 and DNA Methylation in T Cell Development, Function, and Survival. Immunity 15, 763–774 (2001).

2. He, W. et al. Adipose-specific peroxisome proliferator-activated receptor gamma knockout causes insulin resistance in fat and liver but not in muscle. PNAS 100, 15712–15717 (2003).

3. Liston, A. et al. Differentiation of regulatory Foxp3^+^ T cells in the thymic cortex. Proc. Natl Acad. Sci. USA 105, 11903–11908 (2008).

4. Cho, H. et al. Regulation of circadian behaviour and metabolism by REV-ERB-α and REV-ERB-β. Nature 485, 123–127 (2012).

5. Dobin, A. et al. STAR: ultrafast universal RNA-seq aligner. Bioinformatics 29, 15–21 (2012).

6. Trapnell, C. et al. Differential analysis of gene regulation at transcript resolution with RNA-seq. Nat Biotechnol 31, 46–53 (2012).

7. Roberts, A., Pimentel, H., Trapnell, C. & Pachter, L. Identification of novel transcripts in annotated genomes using RNA-Seq. Bioinformatics 27, 2325–2329 (2011).

8. Zhou, Y. et al. Metascape provides a biologist-oriented resource for the analysis of systems-level datasets. Nature Communications 1–10 (2019). doi:10.1038/s41467-019-09234-6

9. Krzywinski, M. et al. Circos: An information aesthetic for comparative genomics. Genome Research 19, 1639–1645 (2009).

